# Reconstruction of compartmentalized genome-scale metabolic models using deep learning for over 800 fungi

**DOI:** 10.1101/2023.08.23.554328

**Authors:** Sandra Castillo, Gopal Peddinti, Peter Blomberg, Paula Jouhten

## Abstract

Eukaryotic metabolism is organized into subcellular compartments enclosed by lipid-membranes. Since most metabolites do not move freely across the membranes, the compartmentalization influences metabolic pathway connectivities. Thus, compartmentalizing the pathways also in genome-scale metabolic models (GEMs) is essential for accurate predictions of eukaryotic cells’ metabolic phenotypes. Compartmentalization has manually been introduced into the model Eukaryote GEMs like for yeast *Saccharomyces cerevisiae*. Non-model organisms’ GEMs can be automatically reconstructed from genome data. However, the existing GEM reconstruction methods do not introduce compartmentalization into the models. To that end, we integrated our novel deep learning protein localization prediction and protein functional annotation into a top-down GEM reconstruction for automatically creating species-specific compartmentalized GEMs. We developed also a universal fungal GEM for top-down reconstruction of species-specifically compartmentalized GEMs and reconstructed models for 834 species. The novel protein localization prediction outperformed the state of the art in classifying proteins into multiple compartments and integrating the enzyme localization predictions to functional annotations improved the top-down model reconstruction. Interestingly, the clustering of the fungal GEMs reconstructed with predicted species-specific enzyme localization resembled more closely the phylogenetic relationships of the species than that of the GEMS without the compartmentalization. Compartmentalization of metabolism in fungi differs from other eukaryotes but the enzyme subcellular localizations vary also among fungal species. The Fungal kingdom encompasses species important for human health, environment, and industrial biotechnology. The reconstructed fungal GEM set offers valuable tools for e.g., predicting metabolic phenotypes of e.g., mushrooms or single-cellular fungi, the roles of eukaryotes in microbial communities, designs optimizing eukaryotic hosts for industrial chemical production, and identifying drug targets against pathogenic fungi. Beyond fungi, the compartmentalized GEM reconstruction method allows combining other universal GEMs for developing cell type specific GEMs for higher eukaryotes including plants, insects, and mammals.

## Introduction

Eukaryotic species manifest varying metabolic enzyme compositions distributed into membrane-enclosed subcellular compartments. Compartmentalization of metabolism in fungi has specific differences to other eukaryotes (e.g., fatty acid beta-oxidation) but the enzyme subcellular localizations vary also among fungal species e.g., pyruvate metabolism (*1*) amino acid biosynthesis and metabolism (*2*) and fatty acid beta-oxidation (*3*) (*4*). Most metabolites do not move freely across the compartment enclosing lipid membranes but their trafficking depends on transport facilitating proteins (*5, 6*). Thus, compartmentalization affects interconnections, redox balancing, and energy provision of metabolic pathways. These processes are determining for the metabolic states the cells may achieve with direct impact in biotechnology (*2*). Cellular metabolic states can be predicted from first-principles of thermodynamics using Genome-scale metabolic models (GEM). GEMs have been manually assembled for model organisms, but simulation-ready GEMs can be automatically reconstructed for any genome-sequenced species (*7*) (*8*) (*9*). However, the current GEM reconstruction methods either omit subcellular compartments (*8*), enforce model organism’s compartmentalization (*10*), or rely on classifying metabolic enzymes in single compartments (*7*). A specific metabolic activity can be distributed in several subcellular compartments. A metabolic activity may have distributed localization due to metabolic isoenzymes with different subcellular localizations (e.g., isocitrate dehydrogenase, alcohol dehydrogenase, malate dehydrogenase, citrate synthase), but also more than 20% of the experimentally validated proteins in Uniprot have been found in more than one subcellular compartment (*11*). Proteins including the metabolic enzymes are sorted into their subcellular compartments in eukaryotic cells according to their amino acid sequence. They may include signal peptides for the sorting in their N-terminal sequence (*12*), as a part of their internal sequence, or in their C-terminal sequence as in the case of peroxisomal targeting (*13*). However, the signal peptides do not carry strict consensus sequences (*14*) and post-translation modifications and protein-protein interactions may influence the protein sorting. Due to the lack of strict consensus sequences, and the complexity of mechanisms involved in the sorting, protein localization prediction is an appealing challenge for machine learning including deep learning (*15*). Deep learning (*16*) is a machine learning approach extracting high-level features progressively from input data through multiple interconnected layers. Deep learning methods have been shown to outperform classical methods for protein localization prediction (*17, 18, 19, 20*). Among them, DeepLoc (*17*) is a state-of-the-art protein localization predictor, a general tool based on a Convolutional Neural Network (CNN) combined with Recurrent Neural Networks (RNN) and an attention layer that attempts to classify proteins into one of ten different subcellular compartments. Most other well-performing methods are also limited to classifying proteins into single compartments. The single-compartment classification is a major limitation preventing the prediction of common multi-localization of proteins in cells (*21*). To that end, we developed a novel deep learning method for multi-localization prediction based on protein sequence, optimized for predicting localization in four important compartments known to enclose major metabolic pathways in eukaryotic cells: mitochondria, peroxisomes, endoplasmic reticulum, and other (i.e., free in cytoplasm and the remaining membrane-enclosed compartments such as nucleus) in which metabolic activities are only starting to be revealed (*22*). Fungal metabolism has implications in human health (*23*) (*24*), food, feed, and beverage fermentation (*25*), and industrial biotechnology (*26*). Fungal microbes are important industrial biotechnology hosts for protein and small molecule production (*27*). Host optimization as well as uncovering the role of fungi in microbial communities like food and beverage fermenting ones, or developing new drugs for fungal infections would benefit from predictive simulations of GEMs (*28*). Fungal genome data, a prerequisite for GEM reconstruction, is accumulating in databases through sequencing efforts. Yet, species-specifically compartmentalized GEMs are lacking for non-conventional yeasts and filamentous fungi. We report here a novel method for automated compartmentalized eukaryotic genome-scale metabolic model reconstruction. It integrates a novel deep learning method for multilabel-multiclass protein subcellular localization prediction, and metabolic enzyme functional annotation using EggNOG orthology resource (*29*) for top-down reconstruction of species-specific models using an algorithm derived from the carveMe method (*9*). Having also developed a universal fungal metabolic model, we can demonstrate the novel compartmentalized eukaryotic genome-scale metabolic model reconstruction method by reconstructing 1582 models for 834 fungal species.

## 1 Results

### 1.1 CarveFungi method for multilabel-multiclass protein sub-cellular localization prediction

An ensemble of 19 deep learning models with different architectures was designed to classify proteins into one or several of four sub-cellular compartments (i.e., mitochondria, peroxisome, endoplasmic reticulum (ER), and a class that included the remaining compartments: Golgi, nucleus, etc., or free in cytoplasm) based on the amino acid sequence. Multi-localization was predicted due to prevalence among eukaryotic proteins (Figure S1)). Details about the deep learning model architectures are provided in the supplementary material. A protein representation of 500 amino acids from the sequence (including the C- and N-terminus) was developed. Protein sequences longer than 500 amino acids were truncated while keeping the two terminal sequences. The protein representation was a combination of three different protein features: seven physicochemical features (mass, hydrophobicity, polarity, pKa, pI, and net charge index of side chains), the protein secondary structure prediction using PsiPred (*30*), and the one-hot representation of the amino acids. These three different protein features were input into the models using different combinations (see supplementary material). The 19 models were trained with a set of 37,000 reviewed proteins obtained from the UniProt database (*11*). Then, a final deep learning model was trained to combine the predictions of the initial 19 models. The final ensemble model was tested using a set of 1426 experimentally validated proteins obtained from the Uniprot database that were not included in the training data. The novel CarveFungi method performed similarly to DeepLoc, a state-of-the-art deep learning method for protein localization prediction (*17*), in the prediction of mitochondrial (with area under the receiver operating characteristic (AUC) curve of 0.88 for DeepLoc and 0.91 for CarveFungi (Figure 1)) and cytoplasmic proteins (with and AUC of 0.86 for DeepLoc and 0.87 for CarveFungi (Figure 1)). However, the CarveFungi method clearly outperformed DeepLoc (*17*) in the prediction of protein localization in the ER (with an AUC of 0.89 for CarveFungi, compared to 0.76 for DeepLoc (Figure 1)) and peroxisomes (with and AUC of 0.84 for CarveFungi, compared to 0.68 for DeepLoc (Figure 1)). Furthermore, in contrast to DeepLoc, the CarveFungi method was designed to assign one or more subcellular localization labels to a single protein.

**Figure 1:**
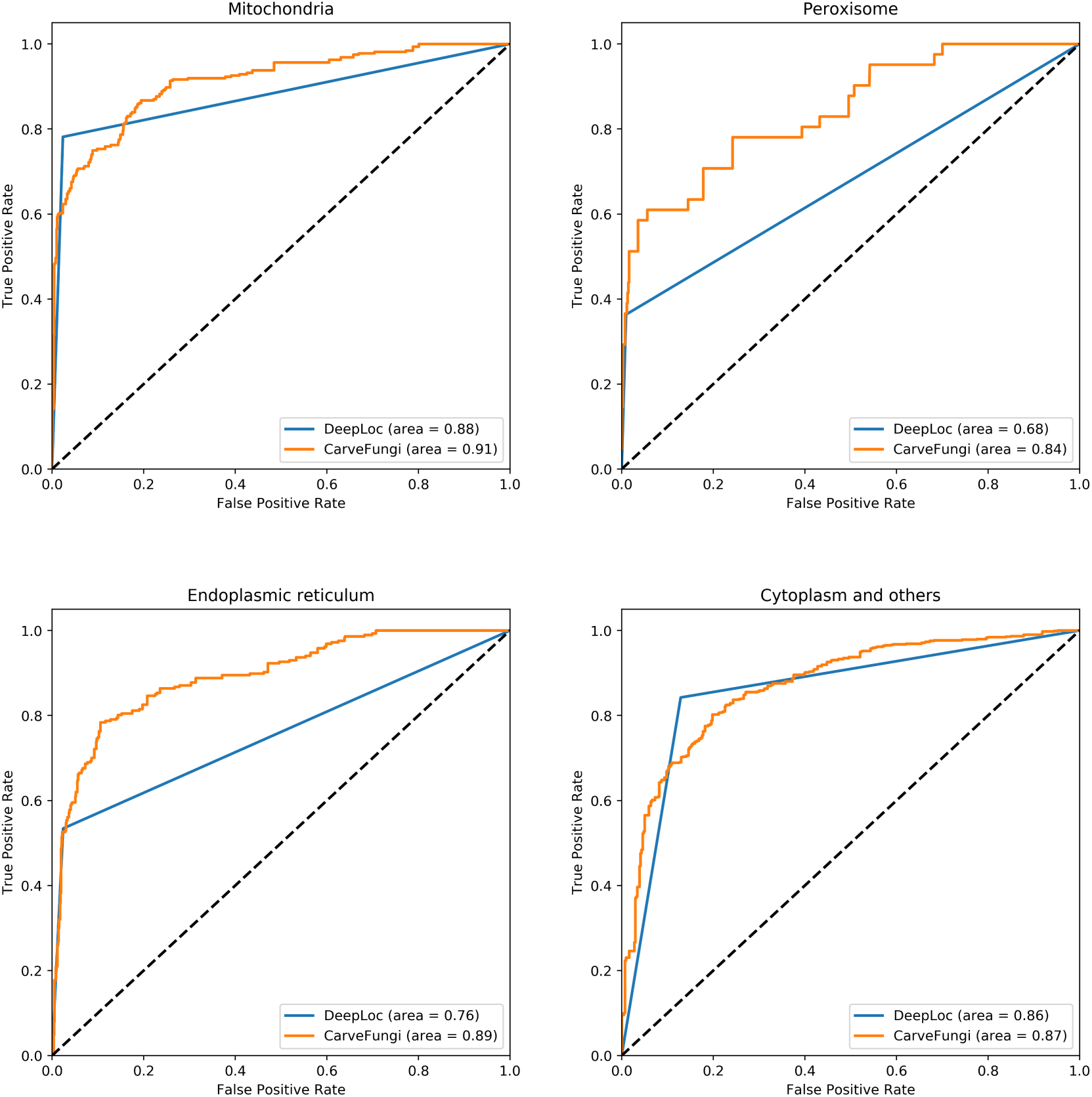
Performance of CarveFungi in protein subcellular localization prediction. Receiver operating characteristic (ROC) curves of the prediction of protein localization in four compartments (i.e., mitochondria, peroxisome, endoplasmic reticulum, and cytoplasm and others (e.g., golgi, and nucleus) with CarveFungi (orange) and DeepLoc (*17*) (blue) methods. When the curve goes along the diagonal (black dashed line) means that the model prediction accuracy is 0.5, same as by chance.

### 1.2 Scoring scheme integrating predicted subcellular localization and functional annotations

Functional annotation of proteins was performed using the orthology resource EggNOG (*29*). The maximum score over all proteins annotated with the particular Enzyme Commission number (EC number) was taken as the evidence that the metabolic enzyme is encoded in the genome. This evidence score of the enzyme being encoded in the genome was integrated with the CarveFungi prediction of metabolic enzyme localizing to the specific subcellular compartment to create a reaction score *S_i,j_* according to Equation 1:

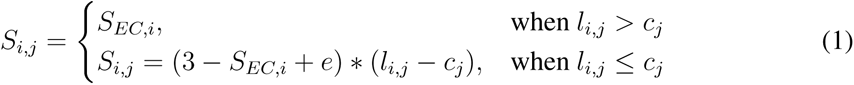

where *S_i,j_* is the score for a metabolic reaction i in subcellular location j, *S_EC,i_* is the highest functional annotation score for enzymes catalyzing reaction i in the genome, *l_i,j_* is the subcellular localization prediction score for a metabolic enzyme catalyzing reaction i in compartment j, *c_j_* a threshold parameter for subcellular location j and e an additive parameter.

The developed scoring of evidence for specifically localized metabolic reactions in cells enables reconstructing compartmentalized GEMs for eukaryotic species. Compartmentalized GEM reconstruction of multiple models could be performed fast, top-down, by removing other than the maximum total score providing reactions from a universal model (*9*).

### 1.3 Creating universal model of fungal metabolism

For fast top-down reconstruction of species-specific compartmentalized GEMs for fungi, a universal model of fungal metabolism was created. Fungal metabolic reactions were retrieved from various sources, including the *S. cerevisiae* consensus GEM (*31*), AYbRAH (*32*) and databases such as KEGG (*33*) and METACYC (*34*). All the reactions to all the compartments except the lipid metabolic reactions for which shared fungal subcellular localizations were assigned. A reaction was then removed from a compartment (i.e., mitochondria, peroxisomes, ER, or other including cytoplasm) in the cases where the reaction was not connected to the rest of the network through a pathway or a metabolite transport reaction reported in a published fungal GEMs (*35*) (*36*) (*37*).The final universal model contained 5063 metabolites and 5602 reactions. They were broadly annotated (e.g., with EC numbers and Kegg (https://www.genome.jp/kegg/) identifiers) to facilitate data integration and extending the model.

### 1.4 Functional annotation of fungal proteins

For fungal compartmentalized genome-scale metabolic model reconstruction all available fungal genomes with gene predictions (n = 1826) were retrieved from NCBI RefSeq (*38*) and Genbank (*39*) databases for functional annotation using the eggNOG-mapper (*29*). While most of the species were annotated with more than ~700 and up to ~1200 enzymatic activities (i.e., EC numbers), some genomes gained lower numbers of annotations (Figure S2). As the small number of enzyme activity annotations indicates insufficient detection of metabolic capabilities, genomes with a number of annotations below 600 were discarded from the model reconstruction. Thus, the model reconstruction was continued with 1582 genomes representing 834 different fungal species.

### 1.5 Compartmentalized model reconstruction for 834 fungal species

Compartmentalized model reconstruction for fungi was initialized by predicting protein sub-cellular localizations for all fungal protein sequences 2. The fungal species differed in the subcellular compartmentalization of the metabolic enzymes 2. Saccharomyces and Candida appeared similar in the numbers of metabolic enzymes in the different subcellular compartments. They differ from filamentous fungi belonging to Aspergilli and Trichoderma. Among Aspergilli major variation in the enzyme compartmentalization was observed.

**Figure 2:**
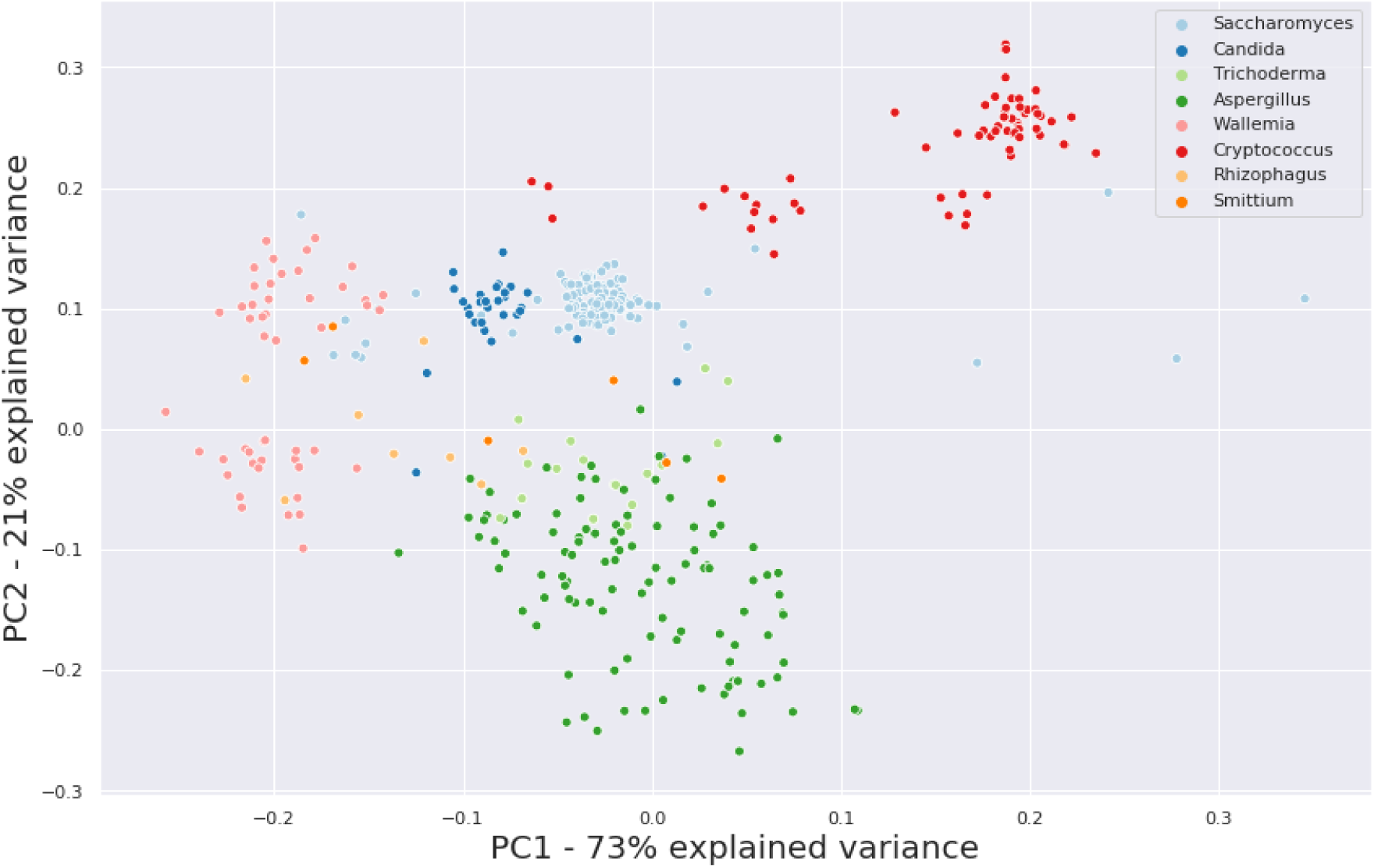
Principal components analysis (PCA) of metabolic enzyme compartmentalization in selected genera. Enzyme compartmentalization is represented by the numbers of metabolic enzymes predicted to be found in the four compartments: mitochondria, peroxisomes, endoplasmic reticulum, and other (i.e., free in cytoplasm and the remaining membrane-enclosed compartments such as nucleus). Each dot represents one species and the colour indicates it’s genus.

Using the scoring scheme developed (Eq. 1) the metabolic enzymes’ scores of the sub-cellular localizations and the functional annotations were used to score the compartmentalized reactions in the fungal universal model. When a reaction was annotated with several metabolic enzyme activities as ECs, the highest EC score was considered to support the reaction belonging to the metabolic capability of the particular species/strain in the specific compartment. Based on the combined functional and localization scoring, species-specific models were extracted from a universal metabolic model using an algorithm derived from carveMe (*9*) (Figure 3).

**Figure 3:**
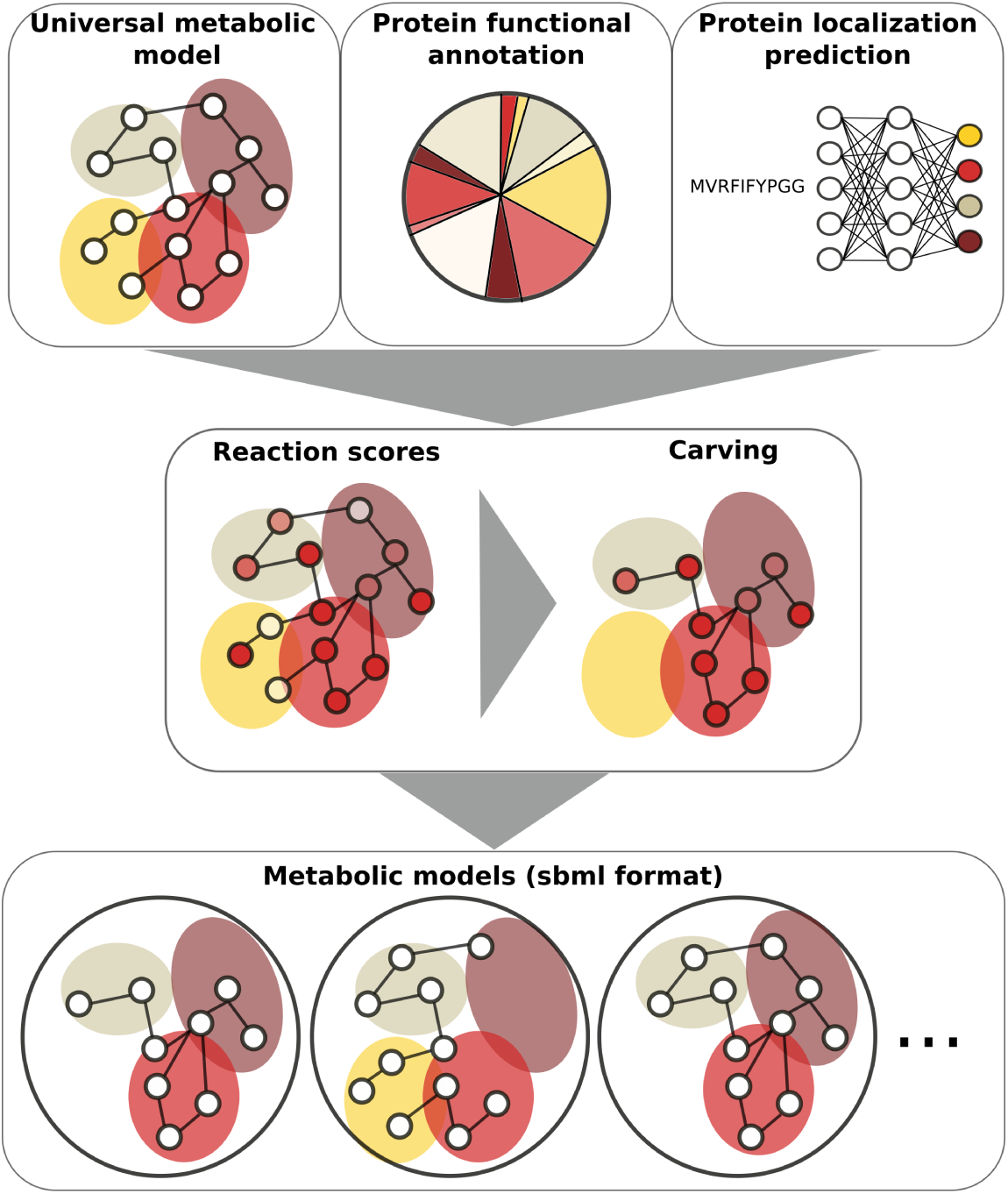
CarveFungi genome-scale metabolic model reconstruction pipeline. CarveFungi performs top-down model reconstruction by scoring the reactions of a universal metabolic model according to protein functional annotation and protein localization prediction. Based on the scores species/strain-specific models are carved out of the universal model using an algorithm derived from carveMe (*9*).

The CarveFungi pipeline (Figure 3) was run twice for each fungal strain: once allowing the organisms to grow on any nutrients available in the universal model, and once by forcing the organisms to grow under aerobic conditions on a defined medium with glucose and ammonium as the sole carbon and nitrogen sources as the minimal media for majority of the species/strains were not known. The algorithm produced one to five models on each run, resulting in two to ten alternative models for each organism. The two to ten models generated in the two runs were then merged in all combinations of models from the first and second rounds of reconstruction, with all nutrients and defined medium, respectively. An ensemble model was then created out of the one to 25 merged models for each strain. A total of 15414 metabolic models were reconstructed, corresponding to 1582 different strains of 834 fungal species. The models were made available at `10.5281/zenodo.7413265`.

### 1.6 Reconstruction accuracy and precision for a model species

The reconstructed ensemble model of *S. cerevisiae* S288C was validated against the metabolic enzymes annotated in *Saccharomyces cerevisiae* Genome Database (SGD) (*40*) and included in the universal fungal model. 243 EC numbers up to four digits (e.g., DNA and RNA ligases) annotated in SGD were out of the scope of the universal model. The obtained accuracy of reconstructing the metabolic capabilities of *S. cerevisiae* was high 0.92 (see table 1).

**Table 1:**
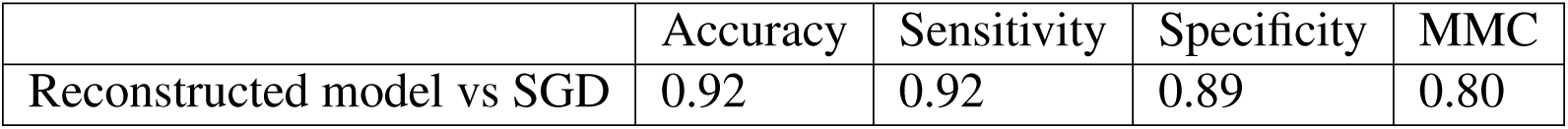
Comparison of our *Saccharomyces cerevisiae* reconstructed model with SGD database. Different metrics to evaluate the comparison of the model reconstruction of *S. cerevisiae* and the SGD database (*40*).

|  | Accuracy | Sensitivity | Specificity | MMC |
| --- | --- | --- | --- | --- |
| Reconstructed model vs SGD | 0.92 | 0.92 | 0.89 | 0.80 |

Since the model reconstruction process could prune the predicted subcellular localizations of metabolic enzymes (Figure S3)), the reaction localizations of the reconstructed ensemble *S. cerevisiae* S228C model were validated against the manually curated yeast consensus model v. 8.3.5 (*31*). For the validation 554 reactions with gene annotations were successfully mapped between the models (see Materials and Methods for details). For the mapped reactions the accuracy obtained was 0.86 for the cytoplasm and others (e.g., golgi and nucleus), 0.96 for the ER, 0.87 for mitochondria and 0.96 for peroxisomes (see table **??**).

The reconstructed *S. cerevisiae* S288C ensemble model was further validated against the yeast consensus model v. 8.3.5 (*31*) by performing essential metabolic gene predictions with the both models with SGD phenotypes of null mutants as the gold standard. 176 metabolic genes were predicted essential using the reconstructed ensemble model simulations. The accuracy of the metabolic gene essentiality prediction was 0.75 and 0.84 for the reconstructed ensemble model and the yeast consensus model, respectively (Table 3). While the automatically reconstructed model performed slightly worse in the accuracy and specificity of the metabolic gene essentiality prediction (table 3), the sensitivity of the prediction was higher using the reconstructed ensemble model (0.69) than by using the yeast consensus model (0.42).

**Table 3:** Comparison of metabolic gene essentiality predictions of the *Saccharomyces cerevisiae* reconstructed models with the yeast consensus model (*31*).

|  | Accuracy | Sensitivity | Specificity | MMC |
| --- | --- | --- | --- | --- |
| Reconstruction of *S.cerevisiae* S288C | 0.75 | 0.69 | 0.76 | 0.38 |
| Yeast consensus model v8.3.5 | 0.84 | 0.42 | 0.91 | 0.34 |

#### 1.6.1 Clustering of phyla by metabolic capabilities was improved by the predicted protein subcellular localizations

Next, it was asked how incorporating predicted enzyme subcellular localizations in the model reconstruction affects the metabolic reaction contents across species and strains. Phylum clustering purity (*41*) was evaluated for models’ reaction content clustering when the models were reconstructed with and without protein subcellular localization scores. Unsupervised clustering (*42*) of the models’ reactions into six clusters was performed (see figure 4). The six clusters represented the six phylum classes of the species and strains for which the models were reconstructed: Ascomycota, Basidiomycota, Mucoromycota, Zoopagomycota, Chytridiomycota and others (i.e., Glomeromycota or Mortierellomycota). The phylum clustering purity was calculated using the equation 2

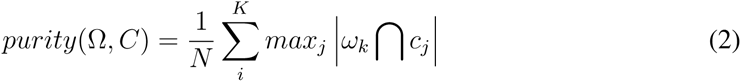

where Ω = {*ω*_1_, *ω*_2_, *ω*_3_, ..} is the set of clusters and *C* == {*c*_1_, *c*_2_, *c*_3_, ..} the set of classes.

The phylum clustering purity was 0.88 and 0.82 for the clustering of the models’ reaction contents when protein subcellular localizations were used and were not used in the model reconstruction, respectively. The phylum clustering purities indicate that the protein subcellular localizations improves the models’ reaction contents as holistically these are expected to follow species relatedness though in specific enzyme activities related species may differ (*43*).

**Figure 4:**
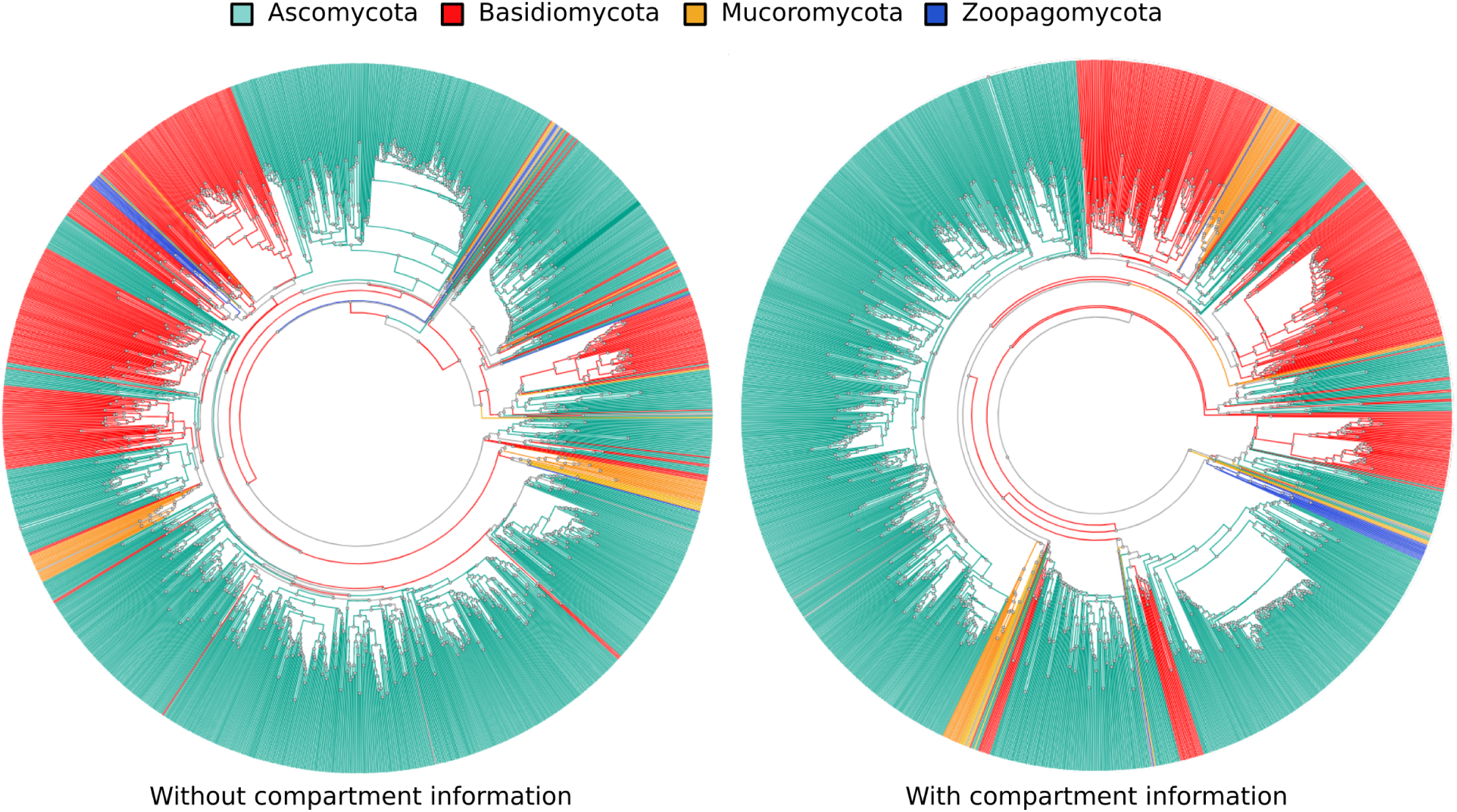
Clustering of the species’ predicted metabolic capabilities with and without subcellular compartmentalization. Clustering of the species’ metabolic reaction ensembles (i.e., out of ECs in the universal fungal metabolic model) predicted without and with subcellular compartmentalization on the left and right, respectively. Species in the clusters are coloured according to the phylum they belong to.

#### 1.6.2 Fungal subcellular protein localization differences reproduced in the reconstructed models

L-lysine biosynthesis is differentially localized among fungi. The first reaction of the pathway, homocitrate synthase (EC 2.3.3.14), has been found localized in cytoplasm in *P. chrysogenum* but in mitochondria and nucleus in *S. cerevisiae* (*44*). The reaction is accordingly localized in the CarveFungi reconstructed models. Another enzyme involved in L-lysine biosynthesis, homoisocitrate dehydrogenase (EC 1.1.1.87), catalyzing the conversion of homoisocitrate to alpha-ketoadipate, has been reported to localize in the cytoplasm in *Schizosaccharomyces pombe* (*45*) but it is mitochondrial in *S. cerevisiae* (*46*) (*47*). This subcellular localization difference is correctly represented in the CarveFungi reconstructed compartmentalized models of the species. Subcellular localization differences among fungi have been reported also in other central metabolic pathways such as in branched-chain amino acid synthesis. *Kluyveromyces lactis* has a single branched-chain amino acid aminotransferase predicted to be localized in mitochondria (*48*) and *S. cerevisiae* has two, a mitochondrial and cytoplasmic version (*49*), allowing functional diversification to synthesis and catabolism (*48*), respectively. The carveFungi *S. cerevisiae* captured the dual localization in *S. cerevisiae*, and predicted mitochondrial localization for the *K. lactis*. Another enzyme involved in the branched-chain amino acid synthesis, alpha-isopropylmalate synthase of L-leucine synthesis route, is mainly mitochondrially localized in *S. cerevisiae* but the minor cytosolic distribution has been found essential (*50*) emphasizing the importance of predicting multi-compartmentalization for modelling (Figure S4)). In *P. pastoris* the distribution is opposite. The majority of the alpha-isopropylmalate synthase activity is found in cytoplasm (*2*) supporting the cytoplasmic prediction by carveFungi. Even a high flux carrying glycolytic key enzyme glycerol-3-phosphate dehydrogenase has been found in different subcellular localizations among fungi. In *S. cerevisiae* there are two isoforms of the enzyme that have different subcellular locations (*51*): cytoplasm and peroxisome. In *Zygosaccharomyces rouxii* glyceraldehyde 3-phosphate dehydrogenase is reported to be cytoplasmic. The reconstructed models accordingly capture also this difference in the reaction subcellular localizations.

## 2 Discussion

In contrast to other automatic genome-scale metabolic model reconstruction tools (*52*) (*8*) (*53*) (*54*) (*55*) (*10*), CarveFungi integrates deep learning for predicting subcellular localizations of metabolic enzymes. Here a deep learning model for predicting fungal protein subcellular localizations was trained but accordingly predictive models could be trained other eukaryotic organisms for modelling their compartmentalized metabolism. The protein features encoding the compartmentalization are complex and often create multi-localization. Protein sequences features signalling for the destination localization may be revealed only by alternative splicing (*56*) not recognizable by conventional predictors. However, the database annotations available for training machine learning models are yet limiting. Here fungal data was sufficient only for the final tuning of the trained models which is not yet optimal for predicting the most subtle compartmentalization differences among fungi. Appropriate compartmentalization is important for accurate eukaryotic metabolic phenotype predictions (*57*). We observed nearly as good metabolic gene essentiality prediction accuracy and specificity and better sensitivity with the automatically reconstructed model with the predicted compartmentalization as can be achieved with a model that has been for years manually curated (*31*). The slightly lower performance in accuracy and specificity may be partially explained by the lack of complex gene-protein-reaction rules in the automatically reconstructed model. The gpr rules build on compositions of metabolic enzyme complexes. The prediction of gprs is likely to rapidly advance by machine learning prediction of protein complexes (*58*) that can be similarly to the protein subcellular localization prediction integrated to metabolic model reconstruction. The compartmentalized metabolic models reconstructed here represent the currently sequenced fungal variety, enriched in Ascomycota and Basidiomycota. The metabolic phenotypes of these fungi predictable using the reconstructed compartmentalized models have important implications in human health, food and beverage fermentations, and biotechnology.

## 3 Materials and Methods

### 3.1 Genome sequence data

Proteome sequences predicted from genome sequencing projects for a total of 1647 fungal assemblies were retrieved from NCBI RefSeq (*38*) and Genbank (*39*) databases. The species and the references to the data are in supplementary material.

### 3.2 Protein functional annotation

The protein sequences were functionally annotated using EggNOG mapper v2 (*29*) with diamond for scoring the sequence similarity, and the EggNOG ortholog database version 5.

### 3.3 Protein subcellular localization prediction

The subcellular localization of proteins was predicted using an ensemble of 19 deep learning models and one extra model that produced the final prediction from the ensemble. Each of the models used a variation of the model parameters and architecture. The input of the models consisted of a matrix where the first dimension of size 500 represented the amino acid sequence, and the second dimension of size seven corresponded to seven different physical and chemical amino acid properties described in Table 4, the secondary structure prediction obtained using PsiPred (*30*), and the one-hot representation of the protein. These inputs were combined in different ways in the deep learning models. The vectors from sequences shorter than 500 amino acids were padded with zeros, and the protein sequences longer than 500 were cut in the middle, leaving the N- and C-terminus intact. The data was pre-processed by clustering with CD-Hit (*59*) and we assigned the clusters to the training (85% or the sequences) and test (15% of the sequences) randomly, to ensure training data sufficiently different from test data. A total of 37000 sequences were used for training the networks and 6229 sequences were used for the tests. This ensured that the results of the tests were not biased by the similarity of the training and test data.

**Table 4:** Physical-chemical amino acid properties. Physico-chemical amino acid properties selected to form the input for the machine learning model.

| Property | Definition |
| --- | --- |
| Mass | Monoisotopic mass of each amino acid divided by the biggest amino acid monoisotopic mass. |
| Polarity | Molecular weight of the whole amino acid divided by the biggest amino acid molecular weight. |
| Hydrophobicity | Physical property that measures how the amino acid is repelled from a mass of water. |
| pKa Value -COOH | Negative of the logarithm of the dissociation constant for the -COOH group. |
| pKa value -NH3+ | Negative of the logarithm of the dissociation constant for the -NH3+ group. |
| pI | pH at the isoelectric point. |
| NCISC | Net charge index of side chains. |

The topology of the neural networks contained one or more bidirectional recurrent layers (*60*) and in some cases, a transformer block (*61*). In some of the models, we included a Dropout layer with probability 0.5 (*62*) for regularization to avoid over-fitting. After those layers, the output was connected to three dense layers starting with 100 neurons and decreasing the number of neurons in each layer. The last dense layer had four neurons corresponding to the four predicted compartments (cytoplasm and others, ER, mitochondria, and peroxisome). The sigmoid function was used as the activation of the last layer of the deep learning models.

The algorithm for parameter optimization used in this work was Adam, an algorithm for first-order gradient-based optimization (*63*). The framework used to create and train the model was Keras (*64*) with Tensorflow (*65*) as a back end. The exact model parameters are found in the supplementary material.

A group of 1426 experimentally validated proteins that were present in the validation set of DeepLoc (*17*) and were not included in the training data was used to perform the final test of the models’ performance (see figure 1). This ensured that the models’ performance was evaluated in a rigorous and fair way.

The thresholds for considering a protein to be in a specific compartment were optimized using part of the test data.

### 3.4 Universal fungal metabolic model

The universal fungal metabolic model was established by combining fungal reactions from several sources, including the *S. cerevisiae* consensus genome-scale metabolic model (*31*), AYbRAH (*32*), KEGG (*33*) and Metacyc (*34*). The lipid and aromatic compound metabolisms were revised and augmented according to the literature by Mäkelä et al. (*66*) and Bugg et al. (*67*). The reactions in the universal model were manually atom balanced and the reaction directionality was assigned through thermodynamic analysis, as in Machado et al. (2018) (*9*). The model was turned simulatable by introducing exchange reactions for uptake of nutrients and excretion of other compounds. To enable growth simulations, *S. cerevisiae* biomass equation from the consensus genome-scale metabolic model v. 7.6 was introduced and augmented with essential vitamins (retinoate, thiamine triphosphate, and biotinate) with small coefficients (0.1 mmol/g CDW). The model was further refined by adjusting flux bounds and introducing missing reactions to allow for biomass formation in minimal medium (i.e., glucose as the carbon source and ammonium as the nitrogen source). The metabolite identifiers were unified and the reactions were annotated with EC numbering and the Kegg identifier when available. The reactions were labeled as either fixed in one of four compartments (i.e., mitochondria, peroxisomes, endoplasmic reticulum, or other, including cytoplasm) or subject to localization assignment through prediction. A significant part of the lipid metabolism was fixed in mitochondria and peroxisomes, as reported by Murayama et al. (*68*), Kohlwein et al. (*69*) and Rambold et al. (*70*).

Transport reactions between compartments were introduced into the universal model based on the various publicly available reconstructed metabolic models ((*71*), (*36*), and (*37*)). Reactions that were not fixed to a specific compartment were cloned in each of the four main compartments (mitochondria, peroxisomes, endoplasmic reticulum and other, including cytoplasm) only if, based on transport reactions obtained from the publicly available reconstructed metabolic models, they were able to carry fluxes under any condition.

### 3.5 Reaction scoring

The reaction scores were calculated as a combination of the EggNog scores from protein functional annotation and the localization probabilities predicted by our deep learning model. Each reaction in the universal model was annotated with one or more EC numbers. The initial score for each reaction was determined by the highest EggNog score of all the sequences annotated with any of the EC numbers in the universal model. These scores were then scaled using a specific factor that was optimized based on the performance of the reconstructed *S. cerevisiae* s288C models against the consensus yeast model v. 8.3.5 (*31*).

When a reaction could potentially occur in multiple compartments, the reaction score was combined with information obtained from protein cellular localization prediction. Only the sequences with an EggNog score higher than the maximum EggNog score minus 50 were considered. The threshold for determining if a protein is present in a particular cellular compartment was optimized using a subset of the test data from the validation set of the compartment prediction method.

The final score for each reaction was not altered if no sequences were associated with the reaction or if at least one of the compartment predictions scores exceeded the threshold for the reaction compartment. Otherwise, it was calculated using the following equation:

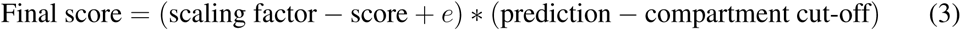

where ‘e’ is a small value (0.00001), the scaling factor is the maximum value of the score after being scaled (3 in this case), the score is the reaction score obtained from the functional annotation (with a range of 0 to 3), ‘prediction’ is a value from 0 to 1 given by the predictive model, and the compartment cut-off was the threshold assigned to each cellular compartment to determine if a reaction is present.

Reactions without gene evidence were assigned a score of minus the scaling factor. Spontaneous, artificial and transport reactions were given a score of 10e−9, and in cases where the EC number was incomplete, the EggNog score was divided by 10.

### 3.6 Top-down model reconstruction

The model reconstruction algorithm was derived from the carveMe method (*9*) which minimizes the number of reactions with low score and maximizes the number of reactions of high score while maintaining the connectivity and functionality of the model. It uses Mixed-integer linear programming (MILP) to find an optimal solution to the problem.

The carveMe method (*9*) was modified to obtain multiple solutions from a pool instead of one. The post-processing of the model to obtain the final model was also changed from the original algorithm. In the original algorithm the small reaction flux in the MILP solution was included as a condition to remove a reaction while we base the inactivation of a reaction only on the binary variables of the MILP solution being less than 0.5.

We combined the five best solutions created from the universal model with open bounds and the five best solutions created from the universal model constrained to grow in minimal media. It gave as a result ten metabolic models and an ensemble of the combination of these ten models written in a single sbml file. To create the ensemble of metabolic models, the presence or absence of each reaction was coded in a string of 25 zeros and ones, and it was concatenated to each reaction name.

For the comparison of our reconstructed *S. cerevisiae* S288C model with the yeast consensus model v. 8.3.5 (*31*), reactions were mapped based on the reaction Kegg id, the compound identifications involved in the reaction and the reaction stoichiometry. We were able to map 554 reactions containing gene annotations.

#### 3.6.1 Metabolic gene essentiality prediction

The metabolic gene essentiality predictions for our reconstructed model for *S. cerevisiae* S288c and the yeast consensus model v. 8.3.5 (*31*) were calculated using the cobrapy function called “single gene deletion”. The results were compared with the list of essential genes obtained from SGD (*40*).

## 4 Author contributions

SC designed and developed the machine learning for protein localization prediction. SC and PJ designed the compartmentalized genome-scale metabolic model reconstruction method and developed the fungal universal model. GP performed the protein functional annotation. PB performed the thermodynamic analysis. SC predicted the fungal protein subcellular localization, and reconstructed and evaluated the genome-scale metabolic models. SC and PJ analysed the results. SC, GP, and PJ wrote the manuscript.

## 5 Data availability

The trained deep learning models for the protein subcellular localization prediction are stored in https://10.5281/zenodo.8272906 and the reconstructed metabolic models can be downloaded at https://10.5281/zenodo.7413265. CarveFungi python code can be found at https://github.com/SandraCastilloPriego/CarveFungi.

## 6 Supplementary material

**Figure S1:**
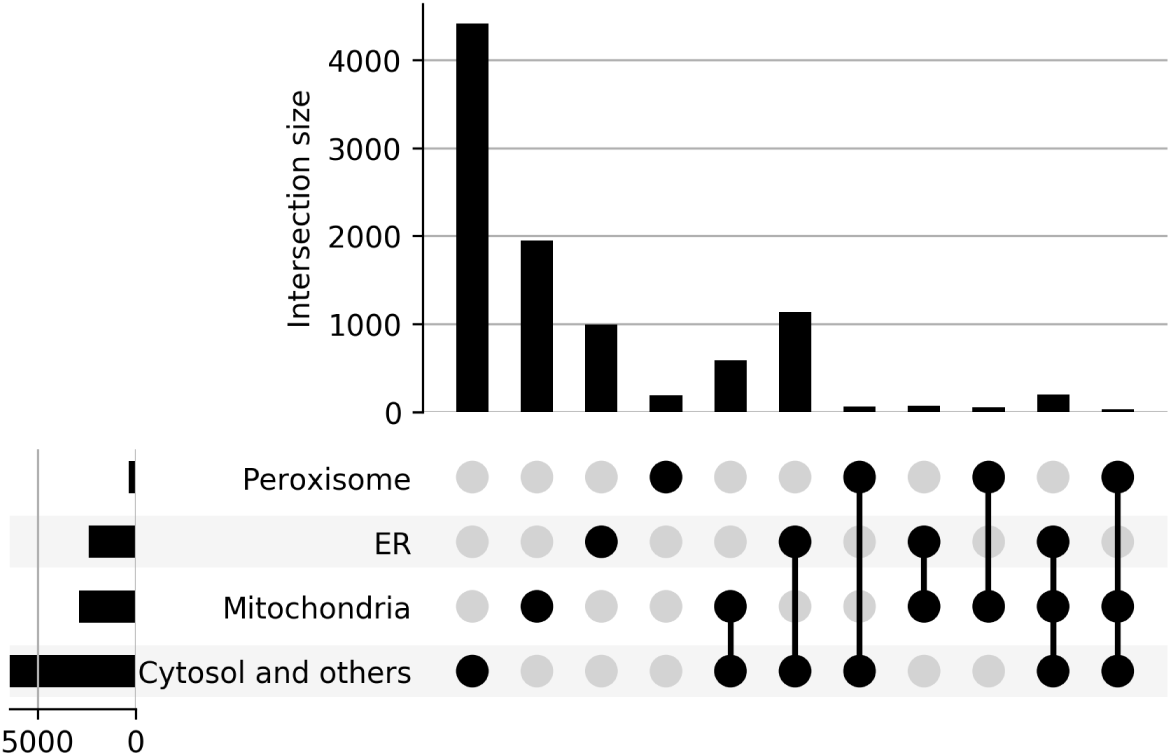
Upset plot of protein subcellular localization. Annotated subcellular compartments of all the curated proteins in Uniprot.

**Figure S2:**
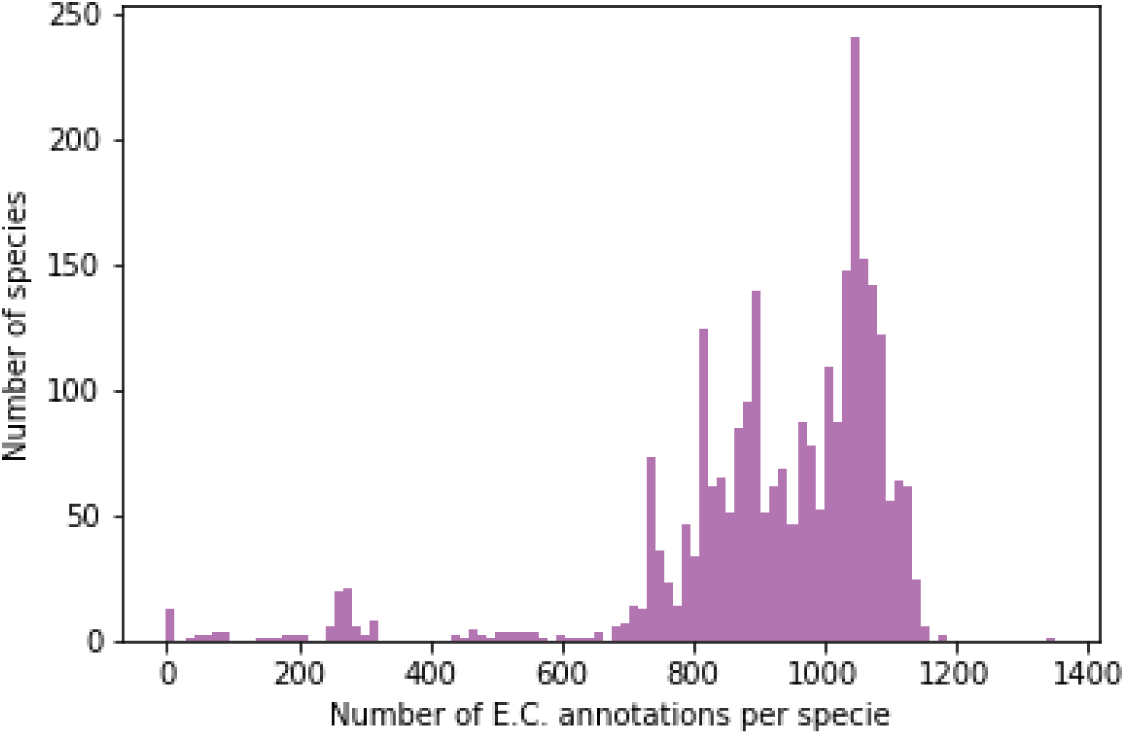
Enzyme commission number (EC) annotations per species. Models were not reconstructed for species with less than 600 EC annotations.

**Figure S3:**
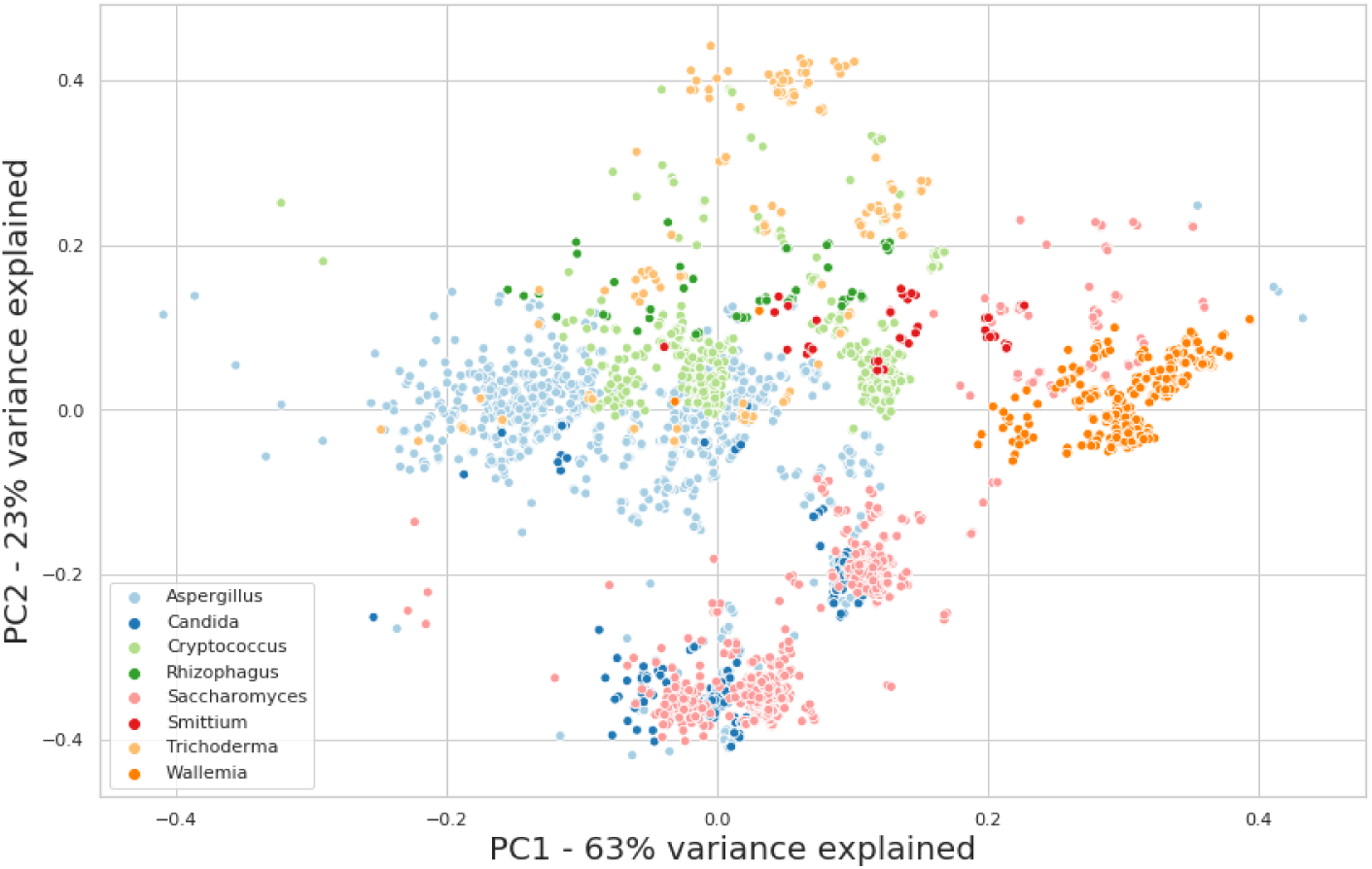
Principal components analysis of the cellular compartment distribution in eight selected species after the CarveFungi compartmentalized genome-scale metabolic model reconstruction.

**Supplementary file 1 — Information on the genome assemblies used in the reconstruction of the compartmentalized genome-scale metabolic models**

**Supplementary file 2 — Model architecture information for the protein subcellular compartment prediction**

**Figure S4:**
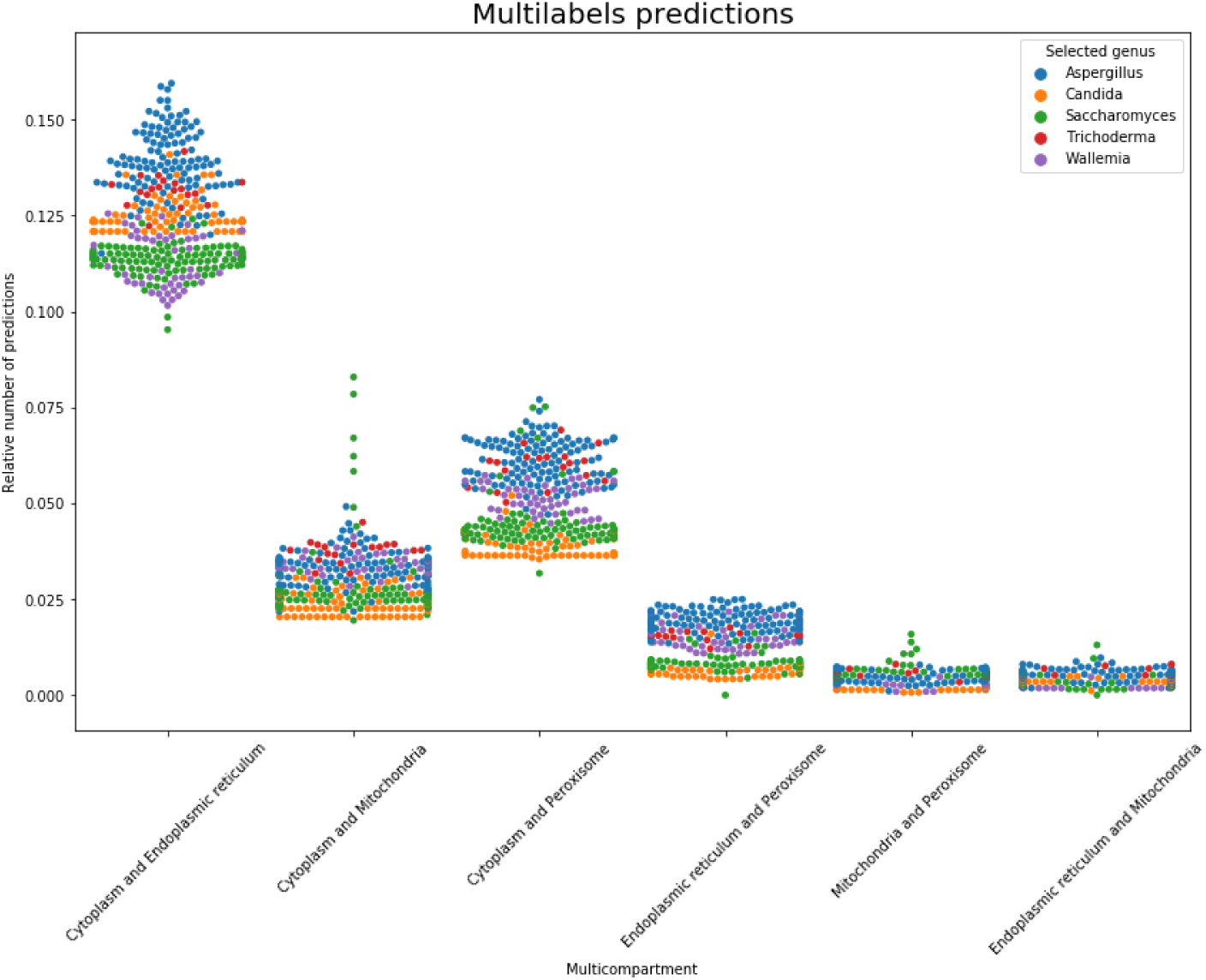
Swarmplot of the multilabel subcellular localization prediction for six selected species.

